# A computational model of typical and impaired reading: the role of visual processing

**DOI:** 10.1101/2021.04.15.440047

**Authors:** Ya-Ning Chang, Stephen Welbourne, Steve Furber, Matthew Lambon Ralph

## Abstract

Computational modelling has served as a powerful tool to advance our understanding of language processes by making theoretical ideas rigorously specified and testable (a form of “ open science” for theory building). In reading research, one of the most influential computational modelling frameworks is the triangle model of reading that characterises the mappings between orthography, phonology and semantics. Currently, most instantiations of the triangle modelling framework start the processes from orthographic levels which abstract away visual processing. Moreover, without visual processing, most models do not provide an opportunity to investigate visual-related dyslexia. To bridge this crucial gap, the present study extended the existing triangle models by implementing an additional visual input. We trained the model to learn to read from visual input without pre-defined orthographic representations. The model was assessed by reading tasks in both intact and after damage (to mimic acquire alexias). The simulation results demonstrated that the model was able to name word and nonwords as well as make lexical decisions. Damage to the visual, phonological or semantic components of the model resulted in the expected reading impairments associated with pure alexia, phonological dyslexia, and surface dyslexia, respectively. The simulation results demonstrated for the first time that both typical and neurologically-impaired reading including both central and peripheral dyslexia could be addressed in this extended triangle model of reading. The findings are consistent with the primary systems account.

---

Computational modelling has served as a powerful tool to advance our understanding of language processes by making theoretical ideas testable and forcing the theories to be rigorously specified. For word reading, there is vigorous development of computational models based on different reading theories (Coltheart, Rastle, Perry, Langdon, & Ziegler, 2001; Harm & Seidenberg, 2004; Norris, 2006; Perry, Ziegler, & Zorzi, 2007; Plaut, McClelland, Seidenberg, & Patterson, 1996; Seidenberg & McClelland, 1989). Of which, the triangle modelling framework has been widely used to characterise the mappings between orthography, phonology and semantics, and their varying roles in different reading tasks (Harm & Seidenberg, 2004; Plaut et al., 1996; Seidenberg & McClelland, 1989). For example, the triangle models of reading can address the division of labour between reading pathways to support effective reading (Harm & Seidenberg, 2004; Plaut et al., 1996), the influences of sequential learning in reading (Chang, Monaghan, & Welbourne, 2019; Ellis & Lambon Ralph, 2000; Monaghan & Ellis, 2010; Zevin & Seidenberg, 2002, 2004), the influences of oral language on learning to read (Chang & Monaghan, 2018; Chang, Taylor, Rastle, & Monaghan, 2020), and the nature of the orthographic written systems (Chang, Welbourne, & Lee, 2016; Yang, McCandliss, Shu, & Zevin, 2009). While these computational models have advanced our understanding of reading, the majority of the models have start from orthographic representations and do not generally take into account the underpinning visual processing. Consequently, the influences of visual-related processing and effects of visual damage have not yet been adequately investigated, and that was the overarching goal of the present study. A brief literature review and the objectives of the present paper are outlined as follow.

Visual processing has been shown to be relevant to a number of psychological effects in visual word recognition, such as letter confusability effects (Bouma, 1971; Gilmore, Hersh, Caramazza, & Griffin, 1979; Loomis, 1982; Townsend, 1971; Van der Heijden, Malhas, & Van den Roovaart, 1984), word-length effects (Balota, Cortese, Sergent-Marshall, Spieler, & Yap, 2004; Weekes, 1997), and letter transposition priming effects (Grainger, Granier, Farioli, Van Assche & van Heuven, 2006; Humphreys, Evett & Quinlan, 1990; Schoonbaert & Grainger, 2004). Of particular relevance here is the relationship between visual processing and word-length effects. Word-length effects refer to that short words are processed more quickly and accurately than long words especially for low-frequency words (Balota et al., 2004; Weekes, 1997). It is originally thought that the word-length effects might be generated purely from sequential processing in the reading system (Coltheart et al., 2001; Rastle, Havelka, Wydell, Coltheart, & Besner, 2009; Weekes, 1997). However, using a variant of the triangle model of reading, Chang et al. (2012c) demonstrated that learning the mappings between visual representations and phonological representations could lead to the model’s sensitivity to letter positions in words, and that was the key to the emergence of word-length effects in the model. If the model started the processes from orthography, the sensitivity to letter positions disappeared (Chang, Furber, & Welbourne, 2012c).

According to the primary systems hypothesis (Patterson & Lambon Ralph, 1999), reading requires interactions of visual, phonological and semantic systems. Damage to different systems is likely to generate different types of dyslexia. For examples, disruption to the phonological system could lead to phonological-related impairments as observed in phonological-deep dyslexia (Crisp, Howard, & Lambon Ralph, 2011; Crisp & Lambon Ralph, 2006), while disruption to the semantic system could lead to semantic-related impairments including surface dyslexia (Hodges & Patterson, 2007; Patterson et al., 2006; Woollams, Lambon Ralph, Plaut, & Patterson, 2007). These are generally termed *central dyslexia* (Shallice & Warrington, 1980). On the other hand, disruption to the visual system could lead to visual-related impairments such as pure alexia (Arguin, Fiset, & Bub, 2002; Behrmann, Plaut, & Nelson, 1998; Damasio & Damasio, 1983; Roberts, Lambon Ralph, & Woollams, 2010) or neglect dyslexia (Primativo, Arduino, De Luca, Daini, & Martelli, 2013; Vallar, Burani, & Arduino, 2010), termed *peripheral dyslexia* (Shallice & Warrington, 1980). Variants of the triangle models of reading have been developed to investigate acquired dyslexia including phonological dyslexia (Plaut et al., 1996; Welbourne & Lambon Ralph, 2005, 2007; Welbourne, Woollams, Crisp, & Lambon Ralph, 2011), surface dyslexia (Plaut et al., 1996; Woollams et al., 2007), deep dyslexia (Plaut & Shallice, 1993) as well as multiple acquired alexia within one framework (Welbourne et al., 2011). However, these models have focused on simulating central dyslexia and do not provide an opportunity to simulate peripheral dyslexia because of the lack of a visual component in the system. There are few notable exceptions of the models that were developed to address visual-related reading deficits (Chang, Furber, & Welbourne, 2012b; Plaut & Behrmann, 2011; Shillcock, Ellison, & Monaghan, 2000). For examples, Plaut and Behrmann (2011) implemented a computational model of visual processing, in which the model was trained to recognise words, faces and houses via hemispheric-specific intermediate connections with topologic constraint on learning. This model demonstrated that general visual processing could support word, face and object recognition, consistent with the view that the higher-level visual processing region (i.e., the ventral occipitotemporal region) serves a general role in processing visual forms (Devlin, Jamison, Gonnerman, & Matthews, 2006; Price & Devlin, 2011; Roberts et al., 2010). Importantly, the model produced graded hemispheric specialisation with words represented more in the left and faces represented more in the right but the patterns were not clean-cut, suggesting pure alexia and prosopagnosia are not entirely pure deficits in words or faces. The predictions by the model are confirmed by subsequent patients studies of pure alexia and prosopagnosia (Behrmann & Plaut, 2014; Rice et al., 2021).

Relatedly, there is a debate on how orthographic representations are formed and stored in the ventral occipitotemporal region of the brain. It could emerge through the interactive processes of learning to read from visual to phonology and semantics (Devlin et al., 2006; Price & Devlin, 2011; Woodhead, Brownsett, Dhanjal, Beckmann, & Wise, 2011). Alternatively, orthographic representations are the stored representations of letter strings and whole words in the orthographic lexicon (Dehaene, Cohen, Sigman, & Vinckier, 2005; Vinckier et al., 2007). The processing principles of the triangle modelling framework are largely in line with the dynamic and interactive view of orthographic representations proposed by Devlin and colleagues. However, for computational simplification, the actual implementation of the models is mostly based on the fixed and pre-defined orthographic representations with only a few exceptions. One previous study demonstrated that the addition of a visual layer allowed the model to develop orthographic representations through learning to read rather than to use pre-defined orthographic representations (Chang et al., 2012c). In all cases, models that have explored visual aspects of reading have not been a full instantiation of the full triangle model, typically omitting semantic representations.

Building on the existing triangle models of reading, this study developed a complete triangle model of reading including visual, orthographic, phonological and semantic processing layers. Specifically, the model started reading processes from visual input and learnt the mappings between visual, phonological and semantic representations; orthographic representations were not pre-defined. The model’s capabilities were assessed on three issues: (a) whether the model could learn the mappings from visual representations to both phonological and semantic representations in which orthographic representations could emerge over the time course of learning and were sensitive to word length; (b) whether the model could perform standard reading tasks including word naming, nonword reading and lexical decision; (c) whether damage to the model could produce the general behavioural patterns of impaired performance observed in patients with the corresponding functional reading deficits. To characterise both typical and impaired reading behaviours in the model, fundamental reading effects were assessed including frequency and consistency in word naming, nonword reading, and imageability and foil type in lexical decision. For the comparison between the intact and impaired models, the same sets of stimuli were used across simulations.

## Simulation 1

Simulation 1 was to construct a computational model of reading based on the general triangle modelling framework (Harm & Seidenberg, 2004; Plaut et al., 1996; Seidenberg & McClelland, 1989; Welbourne & Lambon Ralph, 2007; Welbourne, Woollams, Crisp, & Lambon Ralph, 2011). However, crucially the model started the processing from visual input without pre-defined orthographic representations. The key test was to investigate whether the model could learn the mappings from visual representations to both semantic and phonological representations. Specifically, the performance of the model on reading aloud words and nonwords was evaluated. The model was also tested on the benchmark effect of frequency and consistency.

## Method

### Model architecture

The architecture of the model is shown in Figure 1. The model had two separate pathways for recognising words from visual input: a phonological pathway and a semantic pathway. The visual layer was connected to the OH layer via a hidden layer. The OH layer was equivalent to the orthographic layer in the model, and the representations were learned through the course of training rather than being supplied as inputs. The OH layer was connected to both the phonological and semantic layers via two more hidden layers. Both the phonological and semantic layers were connected to their own set of clean-up units, which helped stabilise the representations. Additionally, there were three context units that provided contextual information for disambiguating homophones. Phonological and semantic units were connected to each other via two hidden layers. With the visual-related layers, the model had a very deep structure. Thus, to effectively regulate the activation of different layers in the model, the control units were added to each layer except input and output layers^1^. These control units received the same inputs as the layer they were connected to and all their outgoing connections were inhibitory. That allowed them to control the activation of all the units in a layer simultaneously.

**Figure 1.**
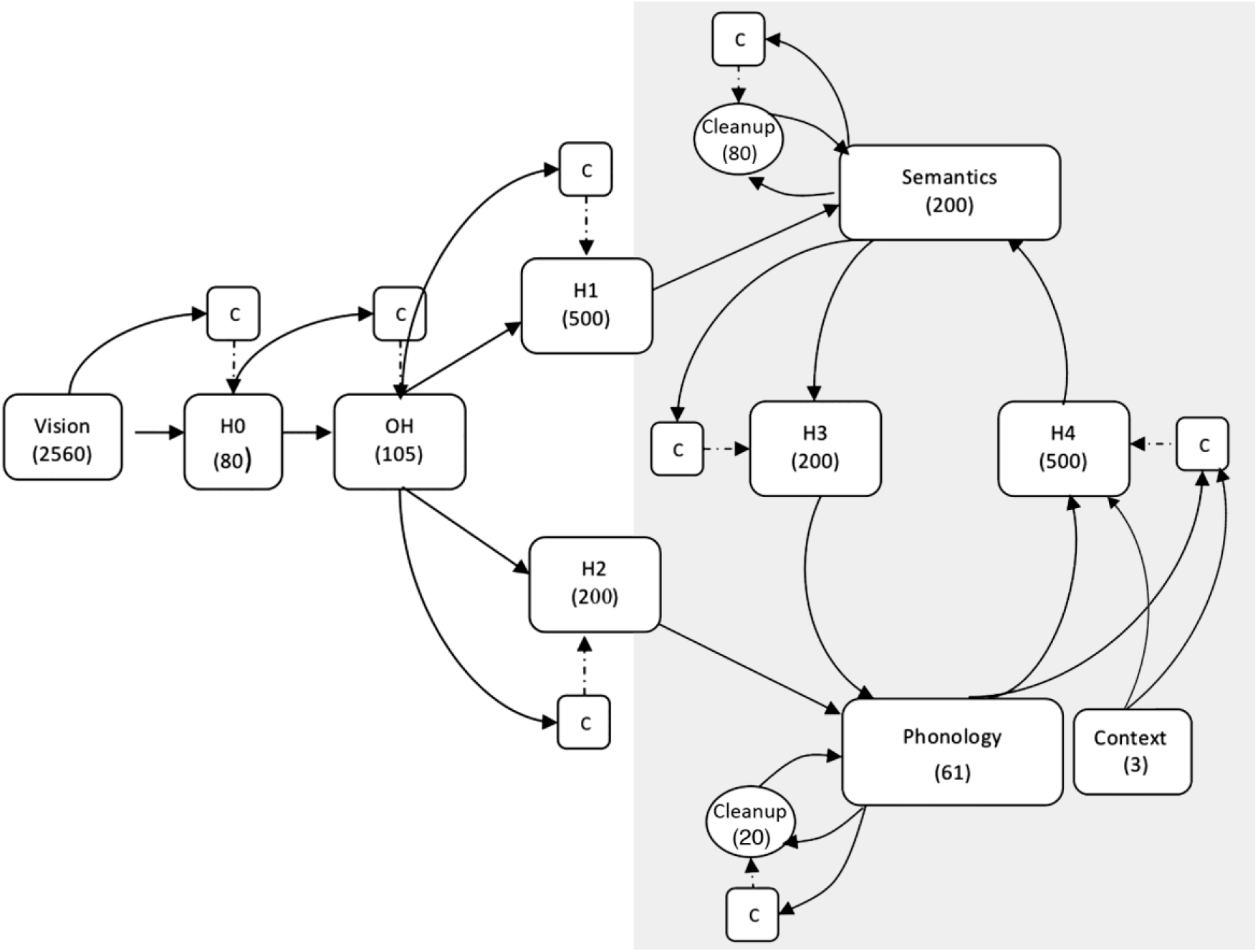
The architecture of the model. It consists of visual, orthographic (OH), phonological and semantic layers connected by five intermediate hidden layers. The two output layers have clean-up units to stabilise the representations. There are local control units for each layer except input and output layers. Each local control unit receives the same inputs as the layer that it controls, and connects to that layer via inhibitory only connections (dashed lines).

### Representations

The training corpus consisted of 2,971 monosyllabic words. For visual input, following Chang et al. (2012c), the model was fed with bitmap images of words in Arial 12-point lower case font. Each word was positioned with its vowel aligned on the central slot of the image. For words that have a second vowel (e.g., boat), the second vowel was placed right next to the first vowel. Ten 16×16 pixel slots were used so there were 2,560 visual units.

The scheme of phonological representations was the same as those used in the Plaut et al. (1996) model. Each word was represented 61 phoneme units which were parsed into onset, vowel and coda clusters of phonemes with specific units used to represent each possible phoneme in each cluster. For the context representations, three context units were used. For non-homophones, the context units were set to zero. Within the same homophone family, different context units were assigned, one for each. The maximum number of meanings corresponding to a given pronunciation in the training corpus is four.

The semantic representations were taken from a semantic space system based on co-occurrence statistics (Chang, Furber, & Welbourne, 2012a). Each semantic representation was composed of 200 semantic units. The key feature of this semantic system is that each semantic vector for an item contains information about the presence and absence of its semantic features. For example, dogs have four legs and they never fly, and both types of information are encoded as one in the semantic vector. Representations derived from this semantic system have validity in reflecting human judgments on semantic category (Chang et al., 2012a).

### Training Procedures

The training procedure was separated into two phases. In phase 1 the links between phonology and semantics were trained (shown in grey in Figure 1) in order to simulate pre-literate language learning in children. In phase 2 the full reading model was trained. All output layers in the model were given a fixed negative bias of −2 to encourage sparse representations.

In phase 1, the phonology-semantics model was subdivided into two parts in which the production model learned the mappings from semantics to phonology, and the comprehension model learned the mappings from phonology to semantics. Both the production and comprehension models were trained on the entire corpus. The probability of each word being presented to the model was determined by its logarithmic frequency. Slightly different learning rates and weight decays were used to train the two models because of the nature of the difficulty of the tasks. The production model was trained with a learning rate of 0.2 and a weight decay of 1E-7. The comprehension model was trained with a learning rate of .05 and weight decay of zero. Each example was presented for six intervals of network time and each interval of time was divided into three ticks. In each presentation, the input pattern was clamped onto the appropriate units six intervals of time. For the last two intervals, the activations of the target units were tested. Error score, the difference between the actual activation and its target activation, was used to calculate weight changes according to the back-propagation through time (BPTT) algorithm (Pearlmutter, 1989, 1995). No error was recorded if the output unit’s activation and its target was within 0.1 of each other. After the separate training, the two models were combined and there was a short period of interleaved training to fine-tune the model.

After the phase 1 training, the weights were loaded into the reading model and frozen so that during reading training, the model was able to utilise the pre-trained knowledge of the mappings between phonology and semantics mimicking children’s oral language skills prior to learning to read. In phase 2, the model was trained on the reading tasks with a learning rate of 0.1, a weight decay of 1E-8 and a momentum of 0.9. Each visual representation was presented for ten intervals of network time (again each interval of time was broken into three time ticks). The model was asked to produce correct phonological and semantic patterns. For the last two intervals, the output activations were compared with their target phonological or semantic representations and errors were computed. No error was computed when the output unit’s activation and target were within 0.001. Again logarithmic frequency was used to determine the probability with which word was presented to the model. To preclude any possibility that simulation result could be generated from one particular set of initial weights, twenty models with different initial weights were trained. Following Bishop (2016), the accuracy rate on regular nonword pronunciations was used to determine the endpoint of training.

### Testing Procedures

The testing procedures for both training phases were exactly the same. The decoding procedure for semantics was based on the Euclidean distances between the activations of the semantic units and each of the semantic representations in the training corpus (Monaghan, Chang, Welbourne, & Brysbaert, 2017; Monaghan, Shillcock, & McDonald, 2004). The semantic representation which was the closest to the activation of the semantic units was taken as the semantic output. If the output was the same as the target representation, it was a correct response. The procedure for the generation of the phonological output was the same as that used in Plaut et al.’s (1996) study. For the vowel units, the most activated vowel unit was selected as output. Onset and coda units were divided into groups of mutually exclusive units and the highest active unit above 0.5 was taken as the output for each group. If no unit was active above 0.5 then the group did not contribute to the output. Finally, if either of the ks ts or ps unit was active along with their components then the order of the components was reversed.

## Results

### Word and nonword reading

At the end of the phase 1 training, the accuracy rates of the production and comprehension model were 99.97% and 99.43% correct respectively. Figure 2 shows the performance of the model over the course of reading training. The average accuracy rates for the model to produce phonological and semantic patterns were 99.3% and 97.4% respectively. Moreover, the model’s generalisation ability was assessed using a set of regular consistent nonwords taken from Glushko’s (1979) study. The pronunciation of a nonword was considered correct if it followed the grapheme to phoneme conversion rules of Venezky (2011), or if it was consistent with the pronunciation of one of the items in the training corpus. The model was able to pronounce 92.6% of the nonwords, which was close to the human performance of 93.8% reported in Glushko (1979). Overall, the simulation results demonstrated that the model was approximately as accurate as human readers at reading words and nonwords.

**Figure 2.**
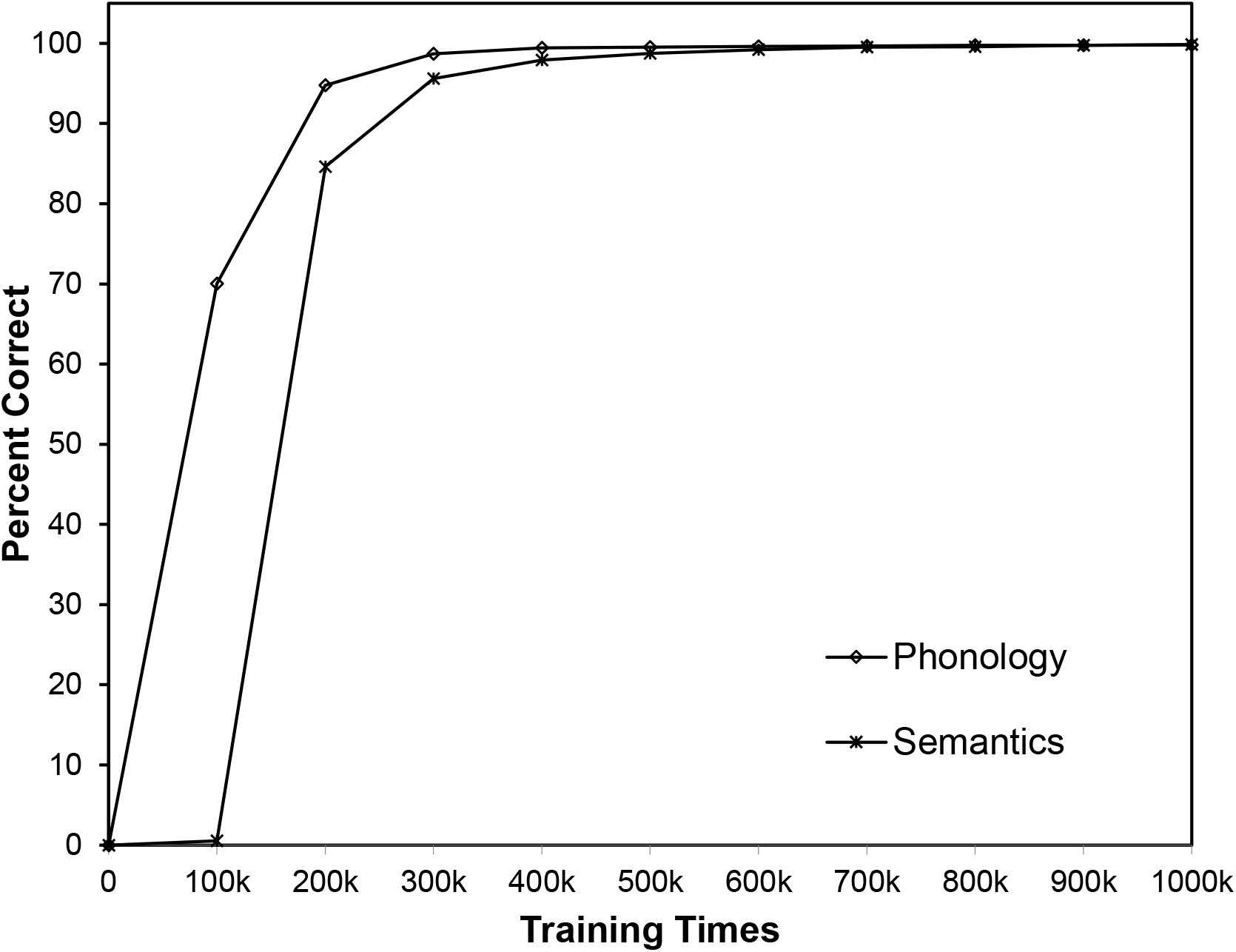
The performance of the model on phonology and semantics during reading training. k indicates1000.

### Frequency and consistency effects on word naming

The model was further verified to see whether it could replicate the benchmark effect of frequency and consistency on word naming (Baron & Strawson, 1976; Jared, McRae, & Seidenberg, 1990; Plaut et al., 1996; Seidenberg & McClelland, 1989; Stanovich & Bauer, 1978; Taraban & McClelland, 1987). Four sets of stimuli taken from Taraban and McClelland (1987) were tested including high-frequency consistent words, low-frequency consistent words, high-frequency inconsistent words, and low-frequency inconsistent words. Each stimulus set consisted of 24 words. Error score was used as an analogy to human reaction times. Figure 3 shows the average phonological error score varying with frequency and consistency. A 2×2 analysis of variance (ANOVA) was performed to analyse the error scores. The main effect of consistency was significant, F(1, 19) = 6.14, *p* < .05, η_p_ = 0.24. The main effect of frequency was also significant, F(1, 19) = 4.95, *p* < .05, η_p_ = 0.21. There was a significant interaction between frequency and consistency, F(1, 19) = 5.55, *p* < .05, η_p_ = 0.23. As expected there was a stronger consistency effect on naming low-frequency words than on high-frequency words, which a pattern similar to that seen in human participants (Taraban and McClelland 1987).

**Figure 3.**
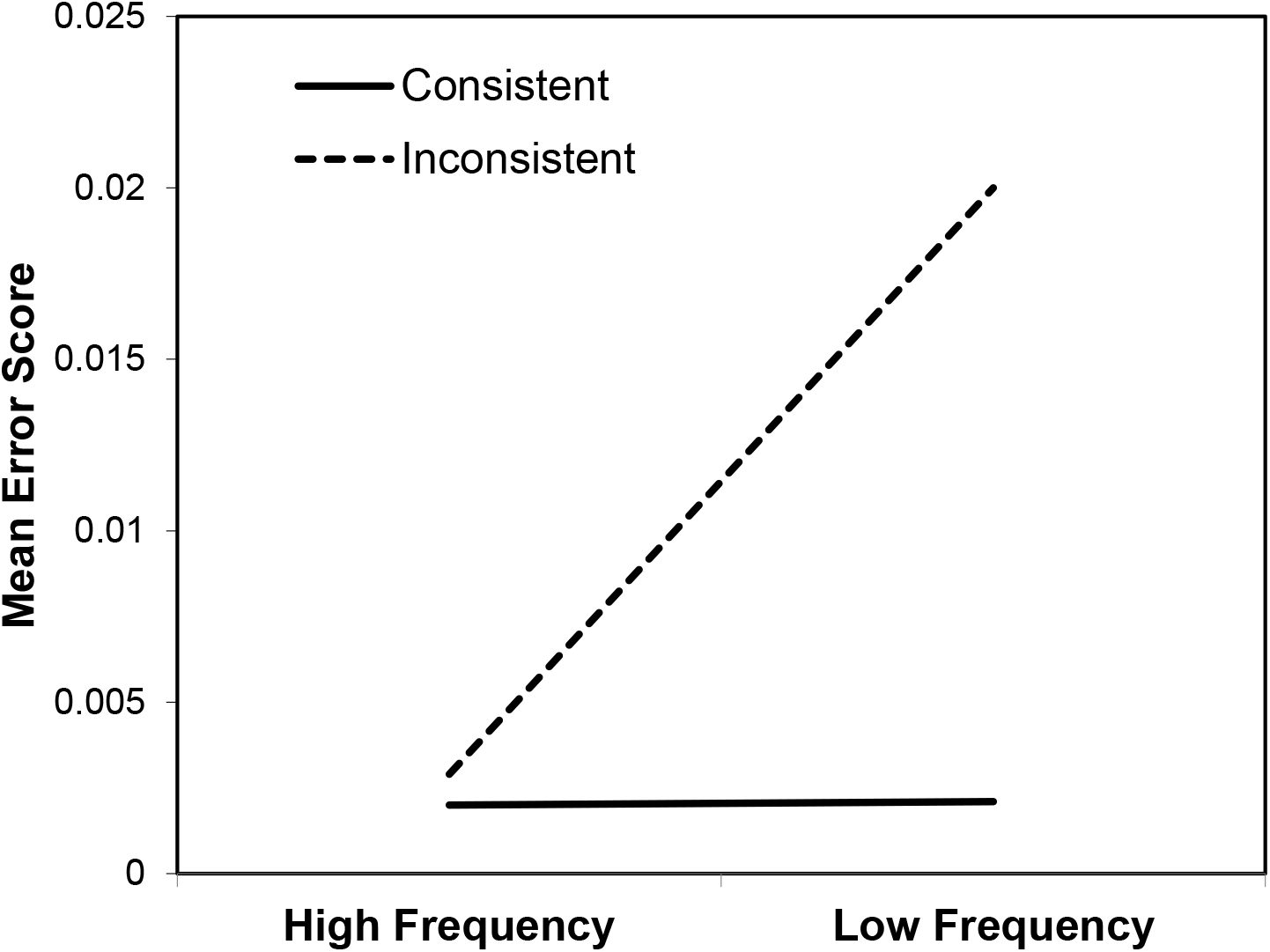
Model performance on the words varying with frequency and consistency

### The emergence of orthographic representations

The simulation results have so far demonstrated that the model could learn to read words and nonwords by training the model on the mappings from visual representations to both phonological and semantic representations. It was critical to further investigate whether the emerged orthographic representations were sensitive to the letter information in words, especially word length, which was the key feature of the triangle model of reading with visual processing as demonstrated in Chang et al.’s (2012c) model. Thus, we extracted the activations from the orthographic layer (i.e., the OH layer) in the model and investigated whether word length was a reliable predictor of the orthographic activations. More specifically, the average activation across all units at the last time tick was computed for all of the words in the training corpus. Then, a linear mixed-effect model analysis was conducted on the average orthographic activations with item and number of models as random effects and with frequency, word length and orthographic neighbourhood as fixed effects. An effect was considered to be significant at *p* < .05 level if its t-value was greater than 1.96 (Baayen, 2008). The results showed that both frequency and orthographic neighbourhood were significant predictors on the average orthographic activations, *β* = −0.117, *t* = −16.44, and *β* = −0.034, *t* = −3.76, respectively. Words with lower frequency scores and fewer neighbours had higher orthographic activations in the model. Critically, word length was also a significant predictor on the average orthographic activations, *β* = 0.153, *t* = 16.72, in which long words had higher orthographic activations than short words in the model. The findings demonstrated, as predicted that the orthographic activations emerged through learning the mappings between visual input and phonology and semantics in the full triangle model of reading were sensitive to lexical variables especially word length.

## Simulation 2

Simulation 1 demonstrated that the model could perform word and nonword reading tasks. We next investigated whether the model could perform lexical decision in which the task was assumed to be supported by the shared processing components in the reading system including, vision, orthography, phonology and semantics.

Plaut (1997) demonstrated that the parallel processing models could perform the lexical decision task based on the measure of polarity in the semantic layer, which is essentially a test of how binary the representations have developed via learning. The idea behind this is that during training, the units are trained to represent target patterns consisting of binary values. Thus when unfamiliar items such as nonwords are presented, the units tend to remain closer to their initial states. To capture this phenomenon, Plaut (1997) used a formula to compute the index of unit binarisation, termed ‘unit polarity’:

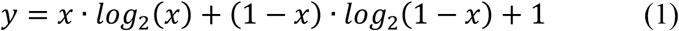

where *x* is the unit activation ranging from 0 to 1 and *y* is the polarity measure.

We adopted the approach used in Plaut (1997) but extended it to include polarity scores not only in the semantic layer but also orthographic and phonological layers in the model because all processing layers should contribute to making lexical decisions, albeit to varying degrees (Plaut, 1997; Seidenberg & McClelland, 1989). Thus, in the present study, polarity scores were integrated across units in the H0, OH, phonological and semantic layers. As both H0 and OH layers were related to the visual-orthographic processes in the model, they were combined as the orthographic processing layer.

### Lexical decision criteria

The average polarity across all units at the three processing layers was first computed separately, and then the grand average polarity score across the processing layers was computed as an integrated measure for making lexical decisions. This procedure was repeated for each time tick in the model (i.e., 30 times ticks in total). The most crucial issue was what criteria should be applied to the polarity measure to allow lexical decisions to be made accurately and quickly? We assumed that the set of criteria should allow the decisions on words or nonwords as quickly as possible when reliable information was obtained. Thus, we developed three criteria for word and nonword decisions: (1) word boundary: If at any tick, the polarity score exceeded the average polarity score for nonwords by more than two standard deviations, then a word decision would be recorded; (2) nonword boundary: If at any tick, the polarity score was more than two standard deviations under the average polarity score for words, then a nonword decision would be recorded; (3) minimum activation: the decision can only be made after the polarity reached an active level of 0.8. The last criterion was to ensure that the model could make a decision based on reliable information. When neither of the criteria was met by the time the last network time tick was reached, the responses were made based on whether their polarities at the last time tick were closest to the average word and nonword polarities. The response latencies for those items were assigned according to their distance to the average polarity. For the top 10% items, the response latencies were 31 ticks, and for the next 10% items, the latencies were 32 ticks, and so on. The slowest responses for the least 10% items were 40 ticks. Note that, this post hoc assessment was under an assumption that the polarities should have reached a boundary if we allowed the model to run longer time intervals though it was computationally too expensive.

### Inverse efficiency

It is worth noting that the cut-off lines for the word and nonwords are arbitrary and could be varied to produce a different speed-accuracy trade-off. To control for these potential differences in a speed-accuracy trade-off, we adopted inverse efficiency as our performance measure. Inverse efficiency is reaction time divided by accuracy, and it is relatively robust to different levels of speed-accuracy trade-off (Roberts et al., 2010; Röder, Kusmierek, Spence, & Schicke, 2007). To illustrate the effectiveness of inverse efficiency, we tested the model on all the 2,971 words in the training set against a set of nonwords consisting of the same number of monosyllabic pseudowords taken from the ARC nonword database (Rastle, Harrington, & Coltheart, 2002). The length of nonwords ranged from three to seven letters. Different cut-off lines were tested including one standard deviation, two standard deviations and three standard deviations. When the cut-off line of one standard division was used, only 0.2% of word responses did not meet the decision criteria but this increased to 27.2% for the cut-off line of two standard deviations and 69.1% for the cut-off line of three standard deviations. The response latencies of those items were assigned according to their distance to average polarity. Figure 4 shows the distributions of accuracy, response time and inverse efficiency for three different cut-off lines produced by the model. The use of different cut-off lines greatly influenced the distribution patterns of response time while the distribution patterns of inverse efficiency remained similar, indicating inverse efficiency was relatively robust to the selection of different cut-off lines. In the following tests, we opted to use the cut-off line of two standard deviations.

**Figure 4.**
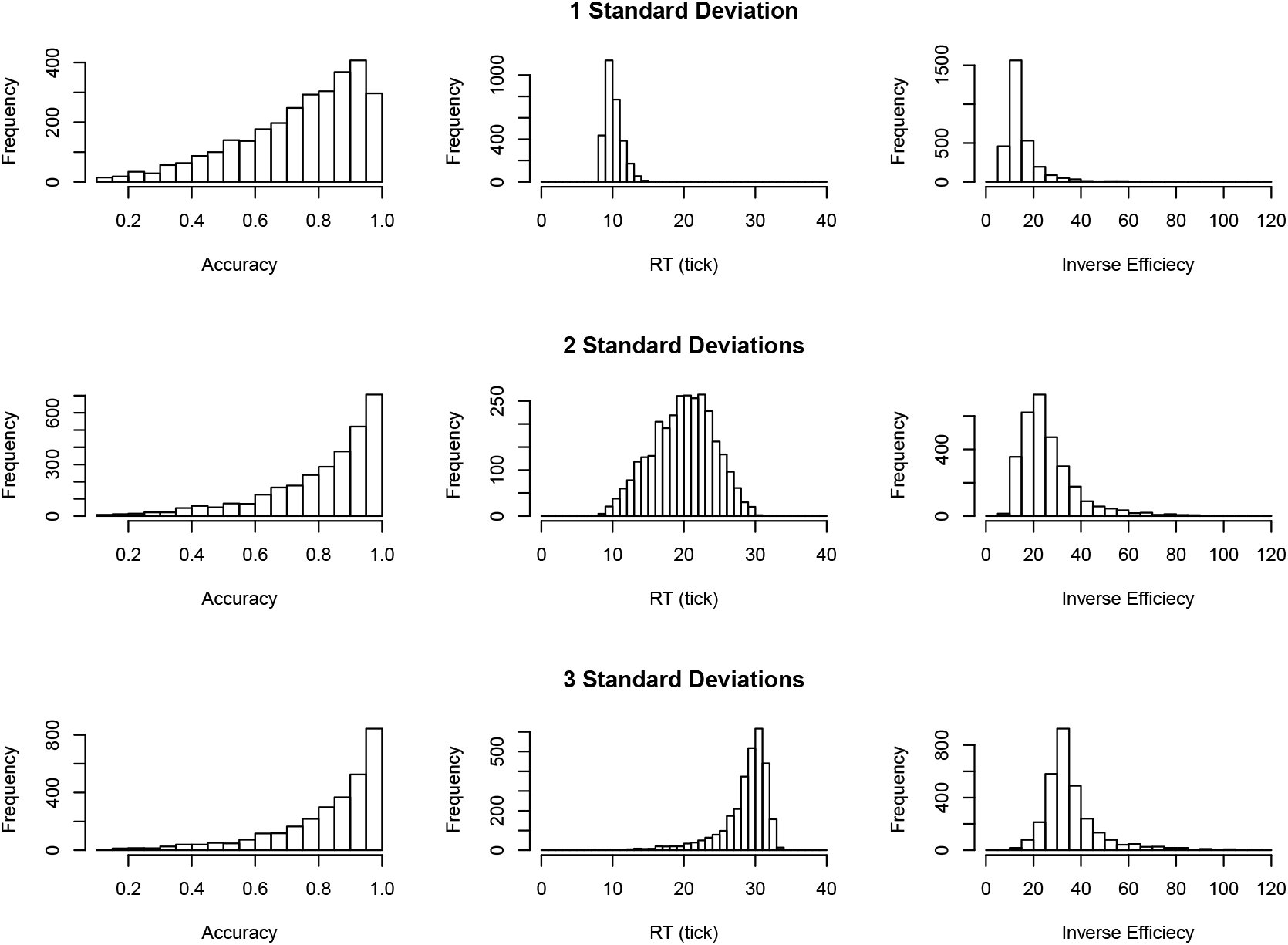
The distributions of accuracy, response time and inverse efficiency for three different cut-off lines (1 standard deviation, 2 standard deviations and 3 standard deviations) produced by the model

### Imageability and foil type on lexical decision

The model was assessed to see whether it could produce a graded imageability effect depending on condition difficulties as observed in Evans et al. (2012) in which the imageability effect was larger when words were tested in the context of pseudohomophones than pseudowords, and it disappeared altogether in the context of consonant strings as in Figure 5 (panel A).

**Figure 5.**
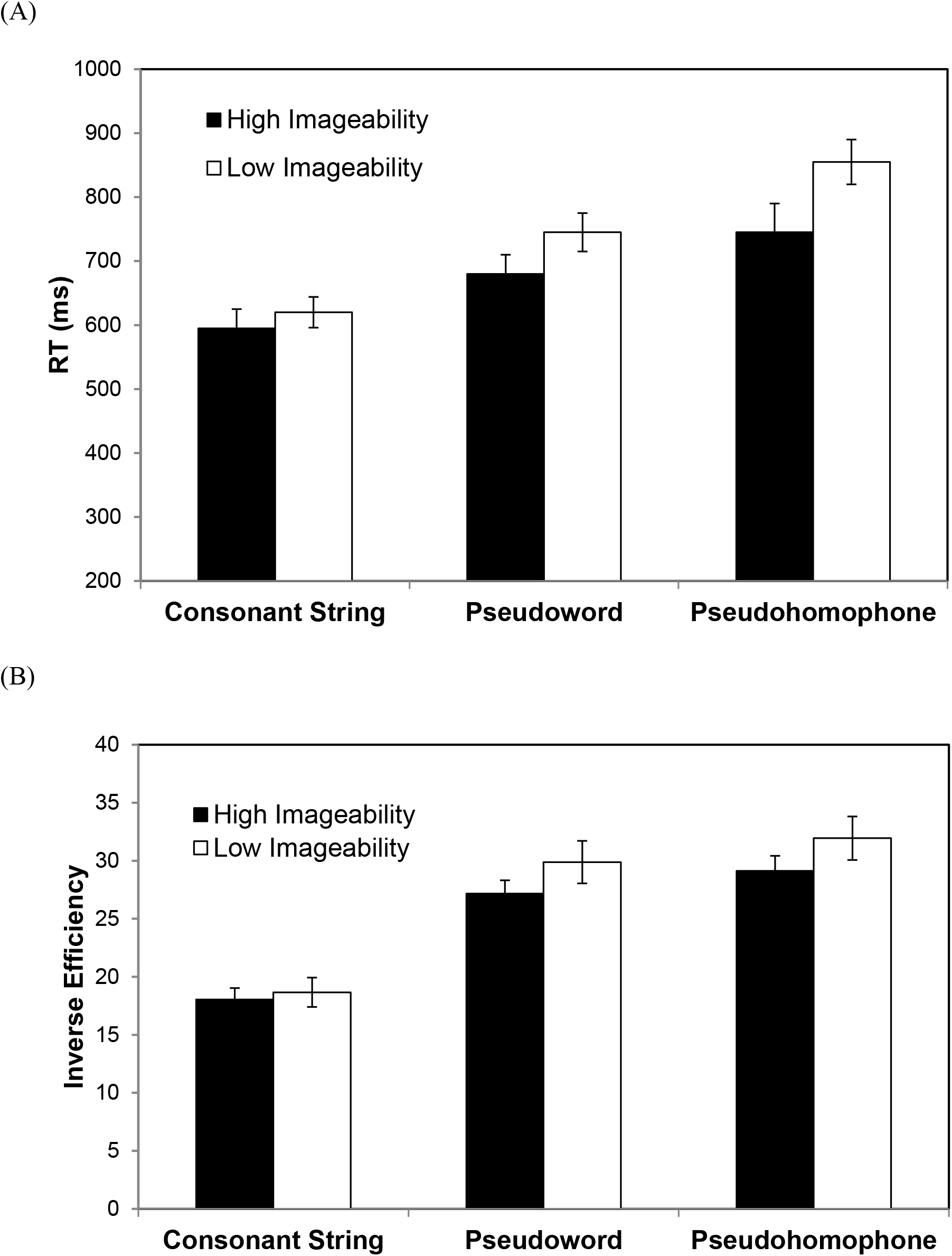
Human data from Evans et al. (2012; panel A) and the simulation data (panel B). Inverse efficiencies for words of high imageability and low imageability in the contexts of consonant strings, pseudowords and pseudohomophones. Error bars represent ± 1SEM.

The model was tested on the stimuli taken from Evans et al. Words that were not in the training corpus and their matched nonword items were removed. The remaining 70 words consisted of 35 high- and 35 low-imageability words along with their matched nonword items. Incorrect items and those with inverse efficiencies farther than three standard deviations from the mean were discarded prior to the analyses. A 2×3 repeated measures ANOVA was used to confirm that the model exhibited a similar pattern of results found in the behavioural data. There were reliable main effects of foil condition, F(2, 38) = 54.09, *p* < .001, η_p_ = 0.74, and imageability, F(1, 19) = 6.16, *p* = .023, η_p_ = 0.245, together with the critical significant interaction between imageability and foil condition, F(2, 38) = 8.21, *p* = .001, η_p_ = 0.302. The simple effect analyses showed that the imageability effect was not significant with consonant strings (*p* > .05). There were significant imageability effects in the context of pseudowords, F(1, 19) = 7.43, *p =* .013, as well as pseudohomophones, F(1, 19) = 7.53, *p* = .013, with the effect size numerically larger for pseudohomophones (η_p_ = 0.284) than for pseudowords (η_p_ = 0.281). The results are shown in Figure 5 along with the comparison of the human data from Evans et al.

## Simulation 3

We next investigated whether damage to different layers would result in patterns of impaired performance similar to those observed in patients with functionally corresponding reading deficits - namely pure alexia, phonological dyslexia and surface dyslexia as predicted by the primary systems hypothesis (Patterson & Lambon Ralph, 1999).

### Word naming and lexical decision in patients with reading deficits

#### Pure alexia

Pure alexia (PA) is a neuropsychological deficit generally caused by lesions in the left ventral occipitotemporal region (Damasio & Damasio, 1983). The PA patients generally show strong word-length effects when naming words, and it is thought by many to result from damage to visual processing (Arguin et al., 2002; Behrmann et al., 1998; Fiset, Arguin, & McCabe, 2006; Roberts et al., 2010). Despite the visual impairment, some PA patients’ word naming performance is still sensitive to lexical variables such as frequency (Behrmann et al., 1998; Johnson & Rayner, 2007; Montant & Behrmann, 2001), regularity (Behrmann et al., 1998), orthographic neighbourhood size (Arguin et al., 2002; Fiset et al., 2006; Montant & Behrmann, 2001), age of acquisition (Cushman & Johnson, 2011) and imageability (Behrmann et al., 1998). For lexical decision, the vast majority of the PA patients’ performance is modulated by frequency (Behrmann et al., 1998; Coslett & Saffran, 1989; Roberts et al., 2010; Staller, Buchanan, Singer, Lappin, & Webb, 1978). Some PA patients’ performance is also sensitive to imageability (Behrmann et al., 1998; Roberts et al., 2010; Staller et al., 1978). According to the partial activation account (Behrmann et al., 1998), it is likely that the lexical-semantic processing could still be partially activated by bottom-up visual stimuli in the PA patients, though that is largely dependent on the severity of the deficit (Roberts et al., 2010).

#### Phonological dyslexia

Patients with phonological dyslexia (PD) are characterized by a relative impairment of nonword reading in the context of better word reading accuracy (Beauvois & Dérouesné, 1979; Patterson & Kay, 1982). Many studies suggest that the functional locus of phonological dyslexia is disturbances to phonological processing because the patients’ reading performance is strongly correlated with their non-reading phonological deficits, and they exhibit the same qualitative performance characteristics on reading and non-reading tasks, including substantial lexicality and imageability effects (Crisp & Lambon Ralph, 2006; Patterson & Marcel, 1992; Rapcsak et al., 2009). With respect to the lexical decision task, this task is generally not a key diagnostic test for phonological dyslexia. Consequently, very few studies have examined it.

#### Surface dyslexia

The vast majority of cases with surface dyslexia come from patients with semantic dementia (SD: Jefferies, Lambon Ralph, Jones, Bateman, & Patterson, 2004; McCarthy & Warrington, 1986; Patterson & Hodges, 1992; Woollams et al., 2007), which is characterised by a progressive degradation of conceptual knowledge associated with atrophy centred on the ventrolateral and polar temporal lobe (Hodges, Patterson, Oxbury, & Funnell, 1992; Nestor, Fryer, & Hodges, 2006). Indeed the link between surface dyslexia and semantic dementia is long-lived and prominent, early descriptions proposed the alternative name of “ semantic dyslexia” for this entity (Shallice & Warrington, 1980; Shallice, Warrington, & McCarthy, 1983). Importantly, the vast majority of SD patients have surface dyslexia (Woollams et al., 2007), showing greater difficulty in naming words with inconsistent spelling-to-sound mappings, particularly for low frequency items, whereas nonword naming ability is relatively preserved. There are a handful of mild SD patients with good reading performance (Blazely, Coltheart, & Casey, 2005; Cipolotti & Warrington, 1995) but often these patients exhibit surface dyslexia as their semantic impairment progresses further (Schwartz, Marin, & Saffran, 1979; Schwartz, Saffran, & Marin, 1980) perhaps reflecting variations in premorbid individual differences in the reliance on semantics for reading aloud (Hoffman, Lambon Ralph, & Woollams, 2015; Woollams et al., 2007; Woollams, Madrid, & Lambon Ralph, 2017).

Regarding lexical decision, numerous studies have shown that the performance of SD patients is significantly poorer than control (Benedet, Patterson, Gomez-Pastor, & Luisa Garcia de la Rocha, 2006; Diesfeldt, 1992; Patterson et al., 2006; Rogers, Lambon Ralph, Hodges, & Patterson, 2004) with some exceptional cases (Blazely et al., 2005; Coltheart, 2004). The discrepancy is likely due to the severity of the impairment (Plaut & Booth, 2006). Notably, the SD patients’ performance in lexical decision is largely dependent on the test condition of stimuli. Diesfeldt (1992) reported that in a visual lexical decision task, a patient, BHJ, performed well when words were tested against consonant strings but had significant difficulty in distinguishing words from more wordlike nonwords such as pseudowords and pseudohomophones. Moreover, in a two-alternative forced-choice paradigm, the SD patients were able to judge orthographically typical words from the relatively atypical nonwords but their performance was significantly impaired in the reverse condition (Rogers et al., 2004).

To simulate the three different types of dyslexia, we damaged the model by lesioning the functional locus/key processing layer in the model. That was, for PA, the emergent-orthographic processing layer was damaged, and for PD and SD, phonological and semantic layers were damaged, respectively. To mimic the recovery process following brain damage, the damaged models were retrained for a period of time (Welbourne & Lambon Ralph, 2005; Welbourne et al., 2011). We tested the patient models on the relevant sets of stimuli to simulate the patients’ characteristic performance in word naming and lexical decision. Specifically, for word naming, all of the three damaged models were tested on frequency and consistency effects in which the stimuli were taken from Taraban and McClelland (1987). For nonword naming, both the PD and SD models were tested on the stimuli taken from Glushko (1979). For lexical decision, the PA and SD models were tested on imageability and foil type effects in which the stimuli were taken from Evans et al. (2012). Note that both word and nonword naming tests were based on accuracy rates as commonly reported in patients’ studies. The lexical decision test was based on inverse efficiency to alleviate the issue of the potential speed-accuracy trade-off demonstrated in Figure 4.

The stimuli used in Simulation 3 were exactly the same as in Simulations 1 and 2, allowing for the comparison between the patient models and the intact model. Relative to the intact model, we predicted that: (a) for word naming, all of the three patient models would show a strong frequency effect. Additionally, the SD model would show a strong consistency effect especially for low-frequency words; (b) for nonword reading, the performance of the PD model would be much impaired; (c) for lexical decision, both the PA model and the SD model would show a strong effect of foil type. Moreover, the PA model would show some sensitivity to imageability while the SD model would not.

## Method

The approach to simulate PA and PD patient types was similar with the only difference being the location of the damage and the amount of retraining required for the model to recover to a stable performance level. For PA damage, 90% of the links connecting to or from the HO layer coupled with 90% of the links into or out of the connected control units were randomly removed. The model was retrained for 400,000 training times with data sampled every 10,000 epochs for the last 50,000 training times. For PD damage, the model was damaged by randomly removing 90% of the links into and out of the phonological layer together with 90% of the links into or out of the connected control units. Again the model was retrained for 400,000 training times with data sampled from every 10,000 training times for the last 50,000 training times. Semantic dementia is unlike the other two deficits as it is the result of a progressive disorder (Hodges et al., 1992). Following Welbourne and colleagues (2007, 2011), we simulated it by repeatedly interleaving very mild damage and retraining. We randomly removed 0.2% of the links into or out of the semantic layer together with the links into or out of the connected control units and then trained the network for one epoch. This process was repeated 400 times.

## Results

The descriptive and statistical results of the PA, PD and SD models, as well as the intact model on the relevant tasks in word naming and lexical decision, were summarised in Tables 1 and 2 accordingly.

**Table 1.** Descriptive results of the intact and the damaged models on the three reading tests in word naming and lexical decision.

|  | Intact model |  | PA model |  | PD model |  | SD model |  |
| --- | --- | --- | --- | --- | --- | --- | --- | --- |
|  | Word Naming | Lexical Decision | Word Naming | Lexical Decision | Word Naming | Lexical Decision | Word Naming | Lexical Decision |
| *Taraban & McClelland (1979)* |  |  |  |  |  |  |  |  |
| High Frequency/Consistent | 100<br>(0) | - | 99.9<br>(0.3) | - | 99.5<br>(0.6) | - | 94.7<br>(0.5) | - |
| High Frequency/Inconsistent | 98.8<br>(2.7) | - | 97.0<br>(2.5) | - | 98.2<br>(2.1) | - | 84.1<br>(0.9) | - |
| Low Frequency/Consistent | 100<br>(0) | - | 99.1<br>(1.2) | - | 94.6<br>(3.2) | - | 93.0<br>(0.6) | - |
| Low Frequency/Inconsistent | 97.9<br>(2.9) | - | 91.5<br>(6.6) | - | 83.1<br>(6.7) | - | 79.7<br>(1.1) | - |
| *Evans et al. (2012)* |  |  |  |  |  |  |  |  |
| Consonant String/High Imageability | - | 18.0<br>(4.4) | - | 27.3<br>(6.3) | - | - | - | 31.2<br>(1.0) |
| Pseudoword/High Imageability | - | 27.2<br>(5.1) | - | 48.4<br>(6.4) | - | - | - | 63.3<br>(5.7) |
| Pseudohomophone/High Imageability | - | 29.1<br>(5.8) | - | 48.8<br>(6.6) | - | - | - | 57.0<br>(1.9) |
| Consonant String/Low Imageability | - | 18.7<br>(5.7) | - | 27.1<br>(7.0) | - | - | - | 29.4<br>(1.2) |
| Pseudoword/Low Imageability | - | 30.0<br>(8.2) | - | 50.5<br>(8.5) | - | - | - | 53.2<br>(2.7) |
| Pseudoword/Low Imageability | - | 31.9<br>(8.4) | - | 52.3<br>(7.8) | - | - | - | 55.6<br>(3.8) |
| *Glushko (1979)* |  |  |  |  |  |  |  |  |
| Nonword | 92.6<br>(4.3) | - | - | - | 57.3<br>(8.8) | - | 78.1<br>(9.2) | - |

Note: Means and standard errors of variables in brackets. Word naming results were based on accuracy rates and lexical decision results were based on inverse efficiency.

**Table 2.** Statistical results of the intact and the damaged models on the two reading tests in word naming and lexical decision

|  | Intact model |  | PA model |  | PD model |  | SD model |  |
| --- | --- | --- | --- | --- | --- | --- | --- | --- |
|  | Word Naming | Lexical Decision | Word Naming | Lexical Decision | Word Naming | Lexical Decision | Word Naming | Lexical Decision |
| *Taraban & McClelland (1979)* |  |  |  |  |  |  |  |  |
| Frequency, F(1, 19) | *n.s.* | - | 23.1<br>*p* < .001 | - | 144.8<br>*p* < .001 | - | 40.1<br>*p* < .001 | - |
| Consistency, F(1, 19) | 11.5<br>*p* < .01 | - | 36.8<br>*p* < .001 | - | 58.3<br>*p* < .001 | - | 246.6<br>*p* < .001 | - |
| Interaction, F(1, 19) | *n.s.* | - | 15.7<br>*p* < .01 | - | 44.4<br>*p* < .001 | - | 10.0<br>*p* < .01 | - |
| *Evans et al. (2012)* |  |  |  |  |  |  |  |  |
| Imageability, F(1, 19) | - | 6.2,<br>*p* < .05 | - | *n.s.* | - | - | - | *n.s.* |
| Foil, F(2, 38) | - | 54.1<br>*p* < .001 | - | 133.4<br>*p* < .001 | - | - | - | 48.5<br>*p* < .001 |
| Interaction, F(2, 38) | - | 8.21<br>*p* < .01 | - | 4.7<br>*p* < .05 | - | - | - | *n.s.* |

Note: *n.s.*: non significant. Word naming results were based on accuracy rates and lexical decision results were based on inverse efficiency.

For word naming accuracy, as expected, all of the three patient models demonstrated a significant frequency effect while the intact model did not. Critically, the SD model demonstrated a strong consistency effect, F(1, 19) = 246.6, *p* < .001, relative to the intact model, F(1, 19) = 11.5, *p* < .01. There was a significant interaction between frequency and consistency in the SD model, F(1, 19) = 10.0, *p* < .01 but not in the intact model (*p* > .05).

For nonword reading, the PD model could only pronounce 57.3% of nonwords correctly as shown in Table 1, which was much worse than that of the intact model 92.6%, demonstrating the hallmark feature of the patients with phonological dyslexia.

For lexical decision, the PA model showed a significant effect of foil type, F(2, 38) = 133.4, *p* < .001, while the imageability effect did not reach significance. Though, there was a significant interaction between imageability and foil type, F(2, 38) = 4.7, *p* <.05. Post hoc analyses showed that in the pseudoword condition, high imageability words (M = 48.4) were processed slightly more efficiently compared to low imageability words (M = 50.0), albeit it did not research significance (*p* = 0.1). Whereas in the pseudohomophone condition, high imageability words (M = 49.9) were processed significantly more efficiently compared to low imageability words (M = 52.4), F(1, 19) = 5.25, *p* < .05. These results demonstrated that the PA model’s lexical decision performance was sensitive to semantics especially for judging words against the more word-like nonwords. Concerning the SD model, the effect of foil type was significant, F(2, 38) = 48.5, *p* < .001 while both the imageability effect and the interaction were not significant.

## General Discussion

The primary aim of this paper was to extend the existing triangle models of reading to addressing visual-related processing in typical and impaired reading by developing a complete model of reading incorporating visual, orthographic, phonological, and semantic components. Simulation 1 demonstrated that the model achieved good performance on naming words and nonwords. The model was also able to produce standard frequency and consistency effects in word naming. Critically, the model developed orthographic representations that were sensitive to word length through learning the mappings between the visual stimulus of a word and its associated phonology and semantics. Simulations 2 demonstrated that the same model could differentiate words from different types of nonwords based on the integrated information from visual-orthographic, phonological and semantic layers. Lastly, when the specific processing layer was damaged such as visual-orthographic, phonological or semantic, the model produced general reading behaviours as observed in patients who have the corresponding functional impairment. Together, the simulation results demonstrated that both typical and impaired reading (i.e., pure alexia, phonological dyslexia and surface dyslexia arsing from semantic dementia) could be simulated in a fully implemented triangle model of reading, consistent with the primary systems hypothesis (Patterson & Lambon Ralph, 1999).

### Visual-orthographic processing in reading

Most previous computational models of reading based on triangle modelling framework have started the processes from orthography by utilising pre-defined orthographic representations. While these models have been successful in addressing reading effects relevant to phonological and semantic processing, they are relatively silent on the role of visual-orthographic processing in reading. Partly, it is because the pre-defined orthographic representations do not generally contain the information about letter positions in words, and partly, there is a lack of visual processing in the system. However, numerous neuroimaging studies have associated the ventral occipitotemporal region with orthographic processing in word reading (Dehaene et al., 2005; Devlin et al., 2006; Price & Devlin, 2011; Vinckier et al., 2007; Woodhead et al., 2011). In particular, Devlin and colleagues have proposed that the ventral occipitotemporal region could act as an interface between visual stimuli and their sound and meaning. The present study provides computational support to this interactive account by demonstrating that orthographic representations could emerge as a consequence of learning mappings between vision, phonology and semantics. Note that Devlin and colleagues’ interactive account of the ventral occipitotemporal region is not limited to reading; rather they argue that the region would engage with any meaningful visual stimuli. Although the present model was only trained with word stimuli, the visual input was designed to be flexible. Future research could extend the present model to accommodate different types of visual stimuli such as objects and faces.

Moreover, several neuroimaging studies in reading have demonstrated that the activations in the ventral occipitotemporal region are moderated by word length during reading (Mechelli, Humphreys, Mayall, Olson, & Price, 2000; Wydell, Vuorinen, Helenius, & Salmelin, 2003), in which long words elicited stronger Blood Oxygen Level Dependent (BOLD) activations than short words, reflecting the demand of the processing. In line with these neuroimaging findings, our analysis of the orthographic representations also demonstrated that long words had higher average orthographic activations than short words in the model. This simulation result corroborated with recent modelling studies (Chen, Lambon Ralph, & Rogers, 2017) suggesting that the average unit activation in neural network modelling could be used as a proxy for the BOLD activations in neuroimaging studies.

### Neurologically-impaired reading

One key advantage of the present model with a visual processing component along with phonological and semantic processing components is that it provides an opportunity for the model to address not only central dyslexia but also peripheral dyslexia. Within the primary systems hypothesis (Patterson & Lambon Ralph, 1999), damage to different processing components in the reading system would cause difficulties in accessing the relevant information as is generally observed in the corresponding types of dyslexic patterns. That is what we have observed in the simulation results generated from the patient models. For example, the simulations of pure alexia demonstrated that the PA model was able to perform word naming and lexical decision despite impaired visual-orthographic processing. Nevertheless, relative to the intact model, the performance of the PA model was more sensitive to frequency in word naming and to foil type in lexical decision while it was less sensitive to imageability in lexical decision. That is consistent with the partial activation account (Behrmann et al., 1998) suggesting that the reading performance of the PA patients may still be supported by the partially activated reading system. Note that the imageability effect in the PA patients’ lexical decision is not universally observed (Behrmann et al., 1998). One possibility could be that, as the modelling result demonstrated, the imageability effect is more likely to be observed when words are tested against more word-like nonwords because the decisions would require semantic access (Evans et al., 2012). Alternatively, one critical factor that affects the PA patients’ performance, which is not simulated here, is the severity of the deficit. In a case-series study, Roberts et al. (2010) demonstrated that in either very mild or too severe cases, the PA patients would not show the imageability effects. This is because for the mild PA patients’ orthographic information could still be reliable for effectively lexical decision while for the severe PA patients’ orthographic activations may not be able to spread to the semantic layer. Future modelling research can be conducted to simulate the severity of reading deficit in PA patients by varying the levels of visual-orthographic damage.

The model with damage to the phonological processing layer produced a poor nonword naming performance while it was still able to read inconsistent words. This is in accordance with the hallmark feature of patients with phonological dyslexia in word reading (Beauvois & Dérouesné, 1979; Patterson & Kay, 1982). The simulations of semantic dementia in word naming showed a strong frequency and consistency effect, consistent with the findings in the SD patients (Jefferies et al., 2004; Patterson et al., 2006). Additionally, the SD model’s nonword reading performance was commensurate with the performance of SD patients (Woollams et al., 2007) and previous simulations of surface dyslexia (Welbourne et al., 2011). Collectively, these results showed a general pattern of surface dyslexic reading, providing support to the view of a strong association between semantic dementia and surface dyslexia (Woollams et al., 2007). Considering the SD patients’ performance in lexical decision, they generally perform well on rejecting consonant strings (Diesfeldt, 1992), presumably the decisions are made largely based on orthographic information. This phenomenon was captured by the SD model, which demonstrated an effect of foil type in lexical decision.

### Limitations and future directions

The present model has demonstrated that both typical and impaired reading could be simulated within the same modelling framework. There are, however, some limitations to this model. First, the model has simulated a range of standard reading effects in word naming and lexical decision. Though, the research in word reading is vast so the specific tasks chosen here are neither comprehensive nor exhaustive. Rather, they were selected to simulate fundamental reading behavioural patterns observed in typical and impaired reading. However, the key processing components involved in reading has been implemented in the model. Thus, future research can extend the present model to simulate various tasks of interest in word reading. Secondly, the model’s lexical decision criteria based on the average word and nonword polarity scores were static rather than dynamic. Although the average word polarity scores could be considered as an expectation for words that participants have already built through their experience with words, at least to some extent, prior to the experiment, the participants could not have built such as an expectation for nonwords (i.e., the average nonword polarity score) before they actually encounter them. Thus, it might be anticipated that the decision criteria used by the participants in the earlier trials of the experiment may be slightly different from those used in the later trials where the decision criteria may gradually become steady. However, in behavioural experiments, there are often practice trials that might help alleviate the issue and build up stable criteria rapidly. Additionally, a growing number of studies have demonstrated the effects of cross-trial sequence on lexical decision (Balota, Aschenbrenner, & Yap, 2016) in which stimulus degradation and lexicality in the previous trial have impacted on the responses to the current stimuli, providing evidence for trial-by-trial adjustments to decision making. However, the underlying mechanism remains to be understood. Within the current modelling framework, it is possible to investigate the issue by dynamically adjusting the current stimulus polarity scores with reference to stimulus polarity scores in the previous trial or to implement a more flexible decision mechanism such as the leaky competing accumulator (Usher & McClelland, 2001). Thirdly, the simulations of different types of acquired dyslexia only consider the patients’ representative reading behaviours. Obviously, there are wide variations within each type of dyslexia (Behrmann et al., 1998; Crisp & Lambon Ralph, 2006; Rice et al., 2021; Roberts et al., 2010; Woollams et al., 2007). The variations could, for instance, result from the severity of reading deficits and premorbid individual differences in reading (Dilkina, McClelland, & Plaut, 2008; Hoffman et al., 2015; Woollams et al., 2017). Numerous studies have shown individuals differ with regard to reading experience and vocabulary knowledge as a consequence of great variations in reading effects of skilled readers (Adelman, Sabatos-DeVito, Marquis, & Estes, 2014; Andrews & Hersch, 2010; Davies, Arnell, Birchenough, Grimmond, & Houlson, 2017; Yap, Balota, Sibley, & Ratcliff, 2012). Individual differences in the degree of semantic reliance during exception word reading could account for different reading patterns observed in patients with semantic dementia (Woollams, Lambon Ralph, Madrid, & Patterson, 2016). Considering both the severity of reading deficits and premorbid individual differences in simulations of acquired dyslexia would be an interesting topic for future investigation.

## Conclusion

To conclude, we developed the complete triangle model of reading incorporating vision, orthography, phonology and semantics to simulate both word naming and lexical decision based on the share processing components. Importantly, the inclusion of visual processing in the model allowed it went beyond the existing reading models to address the emergence of orthographic representations and visual-related reading deficits. The simulation results demonstrated for the first time that both typical and neurologically-impaired reading including pure alexia, phonological dyslexia and surface dyslexia could be simulated in the triangle model of reading.

## Supplementary

Figure S1 shows the performance of the model on phonology and semantics over the course of the training with and without control units. For the model trained with control units, both semantics and phonology learned relatively quickly. The model reached near perfect performance after 600,000 training time. By contrast, when the model was trained without control units, the learning was much slower. This was because the control units allowed the model to learn to manage its own temporal dynamics and to make the most efficient use of the embedded knowledge in the pre-trained connections between semantics and phonology.

**Figure S1.**
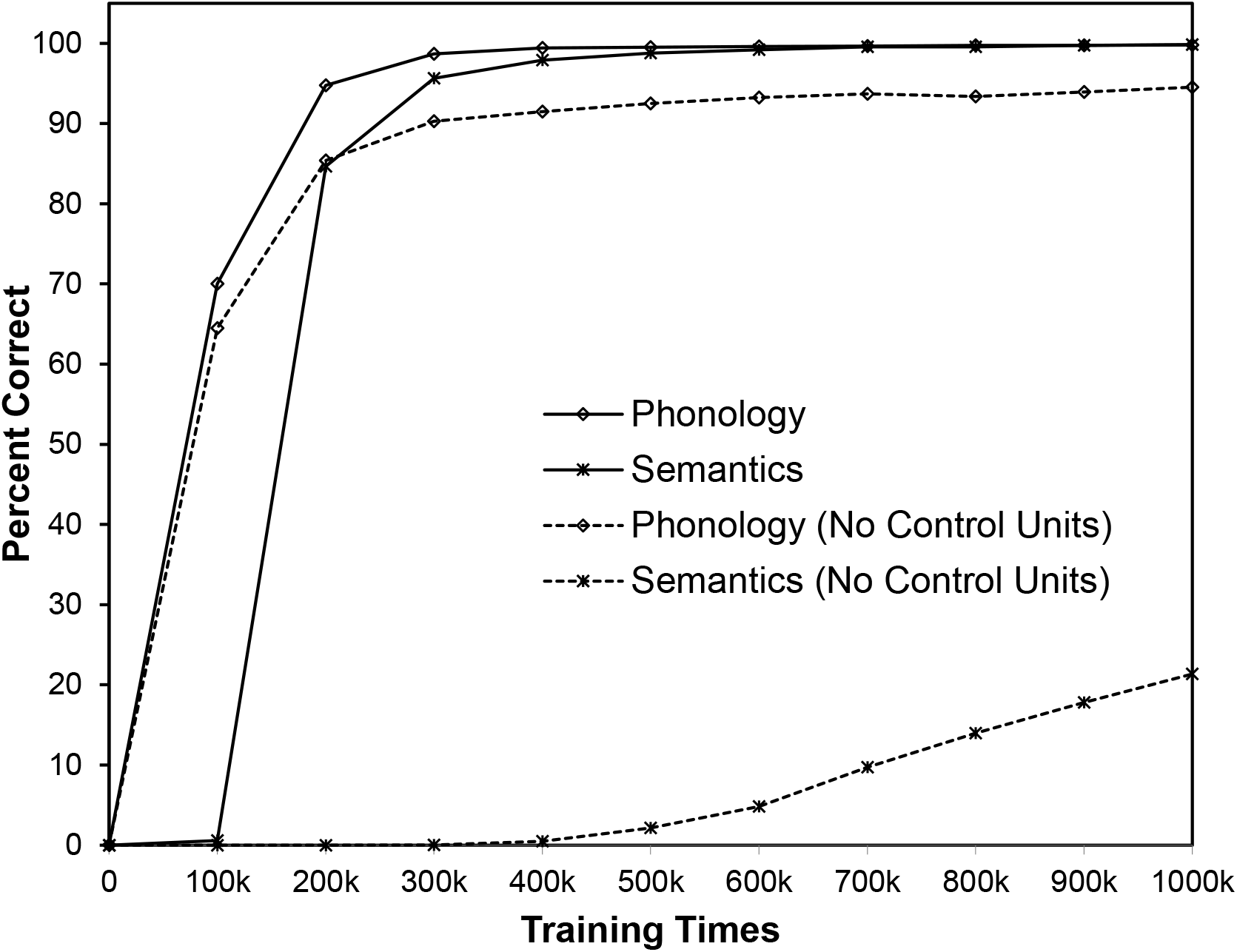
The performance on semantics and phonology in the model trained with and without control units during reading training. k indicates1000.

## Footnotes

1 The control units were important for training the present model with deep recurrent networks, which could help substantially reduce the training time as illustrated in Supplementary.

